# Retrieval-mediated and sleep-based memory consolidation provoke different neural and behavioural markers of memory generalisation

**DOI:** 10.1101/2025.08.15.670466

**Authors:** Hayley B. Caldwell, Kurt Lushington, Alex Chatburn

## Abstract

Across sleep-based and retrieval-mediated consolidation, memories typically become generalised and less dependent on their episodic components for their recollection. However, memory transformations across sleep and retrieval training have not been directly compared. The current study aims to compare how sleep and retrieval training impact the endorsement of semantically similar and different lures, as well as their episodic recollection using the parietal old/new effect on the late positive component (LPC) in subjects’ EEG. Thirty subjects (27F, 18–34, *M*=22.17) attended four sessions where they learnt different sets of 104 object-word pairs and completed one of four 120-minute memory interventions: retrieval training (i.e., cued recall practice), restudy (i.e., pair re-exposure), a nap opportunity, or a wakeful rest. EEG was recorded while subjects were tested on their recognition accuracy in an old/new paradigm with similar- and different-object lures. Our results revealed that retrieval training, but not sleep, lead to greater accuracy for identifying old pairs, but worse similar-lure discrimination. Whilst the parietal old/new effect did not differ between conditions, retrieval had lower LPC amplitudes for similar- than different-object false alarms, whilst restudy demonstrated the opposite. Sleep and wake demonstrated no differences in LPC amplitudes between hits and different false alarm types. Together, our study demonstrates evidence for gist-abstraction across retrieval training, and a task-relevant selective maintenance of episodic details across sleep. These results challenge theories that retrieval training replicates sleep-based consolidation mechanisms, instead acting as a fast route to semanticization regardless of the context.

## 1. Introduction

Memory consolidation allows the brain is to reconstruct relevant previous experiences to readily apply them to current situations (Xue, 2022). Memories are never reconstructed verbatim; instead, our recollection of memories transforms as a function of their semanticization and incorporation into the neocortical landscape. Sleep has long been considered the most optimal state for memory consolidation (Rasch & Born, 2013; Staresina, 2024), although recent evidence has suggested that the same memory transformations can occur across wake, via repeated retrieval training (Antony et al., 2017). A cornerstone of memory transformations across both sleep and retrieval training is that memories are generalised, becoming more dependent on their semantic, neocortical aspects for their recollection than their episodic, hippocampal components (McClelland, 2013; McClelland et al., 1995). Most of this evidence reporting similar influences of sleep-based and retrieval-mediated memory consolidation (e.g., enhanced memory accuracy, semanticization, and shifts to the neocortex) does not compare them in the same paradigm, side-by-side, with adequate control conditions, and where their impacts on memories do not influence each other (Diekelmann et al., 2009; Ferreira et al., 2019; Hebscher et al., 2019; Lifanov et al., 2021; McClelland et al., 1995; Schapiro et al., 2017). To comprehensively understand how memories transform across sleep and retrieval training, it would be informative to directly compare these consolidation types, considering both the behavioural and neural experiences of memory reconstruction thereafter.

Theories explaining the semanticization and gist-extraction of memories across sleep-based consolidation generally convey that memories begin with dual episodic, hippocampal and semantic, neocortical traces. Initially memories are more dependent on the episodic hippocampal traces, leading to more episodically vivid recollections of memories (Born & Wilhelm, 2012; McClelland et al., 1995; Winocur & Moscovitch, 2011). After multiple reactivations of memories that share semantic components but not the contextual episodic components that differentiate them, the semantic components are strengthened, a gist-like representation is constructed, and recalling memories becomes less episodically vivid. Antony et al. (2017) theorised that these same memory transformations occur rapidly across wake, using retrieval training to induce the same type of transformative memory reactivations. They proposed that retrieval training, but not sleep, strongly coactivates related neocortical traces and moderately coactivates competing information to rapidly incorporate memories into the neocortex. Ample evidence exists for these representational transformations and how their reduced veridical recall leads to the behavioural generalisation of memories across retrieval training (Antony et al., 2022; Brodt et al., 2016, 2018; Ferreira et al., 2019; Hebscher et al., 2019; Lifanov et al., 2021; Marin-Garcia et al., 2021) and sleep (Bonnici et al., 2012, 2012; Chatburn et al., 2014; Djonlagic et al., 2009; Gibson et al., 2022; Lau et al., 2011; Lewis & Durrant, 2011; Liu et al., 2024; Takashima et al., 2009; Tompary & Davachi, 2024; Woodard et al., 2007). These studies have illuminated the large-scale semanticization of memories across consolidation. To gain further insight into the experiential impacts of these representational changes, investigating the pitfalls that result from them, and their recollective experiences, will be informative.

Memory generalisation across sleep-based and retrieval-mediated consolidation is often observed to aid inference, category learning, memory for statistical regularities, and paired associations (Ashton et al., 2018, 2021; Bai et al., 2015; Batterink & Paller, 2017; Djonlagic et al., 2009; Ferreira et al., 2019; Javadi et al., 2015; Lerner & Gluck, 2019; Lifanov et al., 2021; Schreiner et al., 2021; Studte et al., 2015; Tompary & Davachi, 2024; Witkowski et al., 2020). However, as gist-like representations are constructed, the episodic components that typically differentiate highly similar memories become less involved in the recollection of those memories. Both naps and full nights of sleep can lead to greater endorsements of critical lures and greater susceptibility to the misinformation effect than periods of wake (Calvillo et al., 2016; Gibson et al., 2022; Huan et al., 2022; Payne et al., 2009; Shaw & Monaghan, 2017). Similarly, retrieval training also leads to greater false alarms and worse lure discrimination than other memory practice conditions (e.g., restudy; Delhaye & Bastin, 2021; Zhuang et al., 2022). Denis et al. (2024) compared retrieval and sleep interventions directly by exposing subjects to a short story before completing either intervention. Post-intervention memory recall revealed that both sleep and retrieval training led to more accurate recall of story components, but a simultaneous greater recall of unpresented, but plausible, information. This design did not control for the effects of general memory practice, which would further aide conclusions about the unique properties of retrieval training. Also, researching a neural measure of episodic recollection to compare memories after sleep and retrieval training would further support their conclusions about gist-extraction across consolidation.

An event-related potential (ERP) is an EEG signal locked to specific events, which captures various ERP components whose mean amplitudes reflect stimulus processing. The Late Positive Component (LPC) is an established ERP component, which occurs approximately 500–800 ms post-stimulus onset. The parietal old/new effect acts upon the LPC over left-lateralised parietal sites, where the LPC amplitude is increased for stimuli that are previously learnt (old), versus unlearnt (new). The parietal old/new effect has been demonstrated to reflect the episodic recollection of memories (Paller et al., 1995; Paller & Kutas, 1992; Rugg & Curran, 2007), which can inversely speak to the extent that memories have undergone semanticization. Several studies have demonstrated that the parietal old/new effect increases as a factor of the amount, quality, and vividness of recollected contextual material, in a variety of modalities, reflecting the reinstatement of a memory’s hippocampal cortical patterns which begins at approximately 500 ms post-stimulus onset (Staresina & Wimber, 2019; Vilberg et al., 2006; Vilberg & Rugg, 2009; Wilding, 2000; Xie et al., 2024; Yick & Wilding, 2008). Whilst studies have found that novel faces and words provoked an increased parietal old/new effect across a period of sleep (Mograss et al., 2008; Palmer et al., 2013), other stimuli related to pre-existing neocortical stores elicited a reduced parietal old/new effect across sleep, relative to wake (Palmer et al., 2013; Zeng et al., 2021), indicating a reduction in episodic recollection suggestive of gist-extraction. Roseburg et al. (2014) found that the parietal old/new effect was increased for material tested twice relative to once. Other studies have shown that repeated retrieval training of schema-congruent and familiar stimuli has instead decreased the parietal old/new effect (Gao et al., 2016; Sweegers et al., 2015). This decreased parietal old/new effect linked with higher behavioural accuracy could have reflected an adaptive reduction of episodic recollection, suggestive of gist-abstraction. This evidence suggests that a reduction in episodic recollection across sleep and retrieval training occurs when stimuli are relevant to existing information and undergo sufficient reactivations. To our knowledge, the parietal old/new effect has not yet been used to measure episodic recollection in sleep and retrieval interventions side-by-side. Doing so will support understandings of how each of these interventions shape recollections and their neural underpinnings.

Another important aspect that may uncover details about the recollection involved in memories that have undergone sleep-based and retrieval-mediated memory consolidation, is the recollection experienced for false alarms. Without intervening consolidation, various studies have shown that false memories evoke similar influences on the LPC with the parietal old/new effect as true memories (Beato et al., 2012; Boldini et al., 2013; Curran & Cleary, 2003; Nessler et al., 2001). One study by Jano et al. (2021) investigated the difference in LPC amplitudes between Deese-Roediger-McDermott true memories and false memories after sleep-based consolidation, and found lower amplitudes for false memories. This was likely due to less episodic information being available for their recollection, as these similar-category (i.e., semantic) lures were formed from reduced episodic fidelity and gist-extraction across sleep. However, we are not aware of any study which has compared recollection in the LPC for hits and false alarms, across retrieval training. Additionally, investigating the episodic recollection involved in false alarms to lures that are semantically similar and different to a target memory could further illuminate how retrieval and sleep impact the experience of false alarms due to gist-extraction (similar lures) versus broader encoding-related failures (different lures).

Taken together, if sleep-based and retrieval-mediated consolidation both involve similar memory generalisation and gist-extraction processes, then both should lead to greater accuracy than control conditions, but also less discrimination of similar lures. Additionally, both conditions should experience a reduced LPC amplitude for correctly identified old, relative to new, stimuli. Lastly, we predict that retrieval training and sleep will lead to reduced LPC amplitudes for similar lures compared to different lures. In the current study, subjects experienced four conditions in which they learnt object-word pairs. Each session, subjects underwent one of four 120-minute memory interventions (retrieval, restudy, sleep, or wake). EEG was used to estimate their parietal old/new effect while they were tested on their memory in an old/new paradigm, with similar- and different-object lures.

## 2. Methods

### 2.1. Subjects

Thirty individuals (27 women, 1 non-binary, 2 men) aged 18–34 years (*M*=22.17, *SD*=4.21) participated in the study. Twenty-five completed all four conditions (retrieval, restudy, sleep, and wake), and five subjects completed 1–2 conditions before withdrawing. All collected data was used in analysis. Subjects were right-handed, fluent English speakers, had normal or corrected vision, and self-reported no hearing or sleep problems. Subjects also reported on interview and questionnaire no diagnosed psychiatric or sleep impairments, and were not taking any medications or recreational drugs impacting EEG in the last 6 months (Massand & Bowler, 2013; Nicholls et al., 2013; Weymar et al., 2010). Subjects were recruited via flyers posted around the university campuses, social media, and an online subject recruitment system. Subjects who completed all four conditions received a $200 honorarium. The University of South Australia Human Research Ethics Committee granted ethics approval for this project (protocol no. 205130).

### 2.2. Materials

#### 2.2.1. Pittsburgh Sleep Quality Index (PSQI)

Sleep habits across the last month were screened using the PSQI (Buysse et al., 1989). The PSQI contains 19 items which are used to generate a total score where scores >5 indicate significant sleep difficulties, which was used as a cutoff for eligibility. The PSQI has demonstrated adequate internal consistency and high test-retest reliability (Backhaus et al., 2002; Buysse et al., 1989; Carpenter & Andrykowski, 1998).

#### 2.2.2. Morningness-Eveningness Questionnaire (MEQ)

The MEQ was used to screen out subjects with extreme circadian types (Horne & Östberg, 1976). The MEQ is a self-report questionnaire involving 19 multiple choice questions on sleep/wake habits. Subjects with scores indicating extreme morningness or eveningness (≤30 or ≥70) were excluded. The MEQ’s items have previously demonstrated a high internal consistency and high test-retest reliability (Li et al., 2011; Pišljar et al., 2019).

#### 2.2.3. Flinders Handedness Survey (FLANDERS)

Handedness was screened for using the FLANDERS (Nicholls et al., 2013). Subjects were instructed to select their hand preference for each of the 10 tasks (e.g., drawing). Selecting “right” corresponds to +1 to their overall score, “left” is –1, and “mixed” is 0. Overall scores between –10 to –5 indicate left-handedness, –4 to +4 indicate mixed-handedness, and +5 to +10 indicate right-handedness. Subjects with scores <5 were excluded. The Flanders has displayed high internal consistency between its items (Nicholls et al., 2013).

#### 2.2.4. Karolinska Sleepiness Scale (KSS)

The KSS was used to control for sleepiness in analyses, due to the reported effects of sleepiness on task performance (Åkerstedt et al., 2014; Kaida et al., 2007; Smith et al., 2002). The KSS is a single-question self-report measure where subjects rate their current sleepiness on a scale from 1–10. Answers range from “1 – extremely alert” to “10 – extremely sleepy, cannot keep awake” (Åkerstedt & Gillberg, 1990). The KSS has demonstrated higher predictive validity for various measures of alertness than any other sleepiness measure, including reaction time (*r*=.57), number of lapses (*r*=.56), and power spectral density in both alpha (*r*=.40) and theta (*r*=.38) bands (Kaida et al., 2006).

#### 2.2.5. EEG

EEG was collected using a 64-channel passive BrainCap with Ag/AgCl electrodes positioned according to the modified 10–20 system. Electrooculographic activity was measured via two electrodes embedded in the cap, positioned approximately 1cm diagonally away from the outer canthus of each eye. Electromyographic activity was measured via three electrodes (two placed on the skin above each subglottic muscle, and one placed between the subglottic muscles), and electrocardiographic activity was measured by one electrode placed above the heart. Two BrainAmp amplifiers (DC system, sampling rate 500 Hz) amplified the signals, while the BrainVision Recorder software recorded the output. Impedances were kept below 10kΩ throughout all experimental phases.

#### 2.2.6. Stimuli

##### 2.2.6.1. Objects

104 object images were taken from the Mnemonic Similarity Task (MST) from sets 2–4 and with false alarm rates at or below 33.33% i.e., chance level (Stark et al., 2019). Each object in the learning set was semantically similar to another object within the set (e.g., a bouquet of red tulips and a bouquet of roses).

##### 2.2.6.2. Words

Word stimuli were taken from an updated version of the Affective Norms for English Words (ANEW; Warriner et al., 2013). This dataset contains 13915 English words, optimised for use in paired associate tasks, rated on frequency from 0–1000 and 1–9 on valence, arousal, and dominance. This study randomly selected 416 monosyllabic nouns (104 per condition), with valence and arousal ratings between 3.5–6.5. These words were recorded, using the voice of a native English speaker with an Australian accent, for their auditory presentation. Each recording was approximately 500–1000 ms long. Words were presented at approximately 70 dB.

### 2.3. Procedure

Eligible subjects were screened using the PSQI, MEQ, and FLANDERS. Subjects then participated in four conditions, each of 4.5 hours (12:00–16:30; see Figure 1), separated by at least 48 hours. The night before each condition, subjects were asked to wake up one hour earlier than usual to increase the likelihood of entering NREM sleep in an afternoon nap, in the sleep condition. Subjects were seated in front of a dedicated computer and were fitted with an EEG cap. Subjects were then given details about the memory task performed across the rest of the session.

**Fig. 1.**
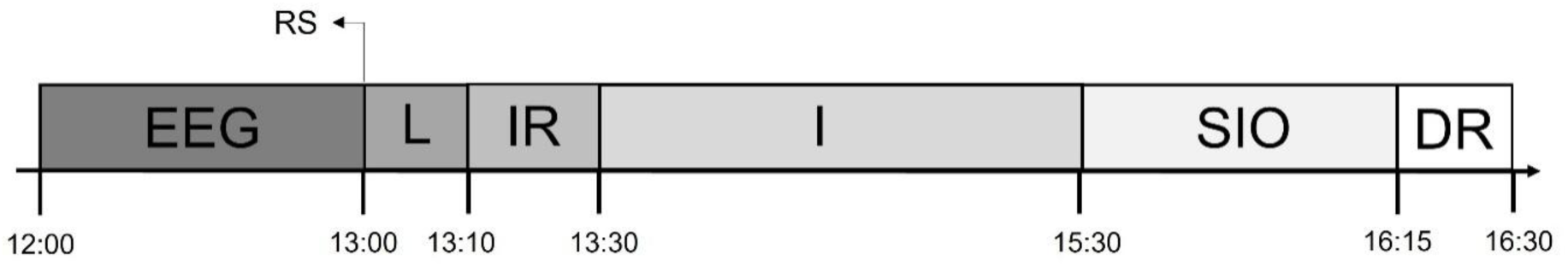
Procedure timeline. EEG = Setup EEG, RS = Resting State, L = Learning, IR = Immediate Recognition, I = Intervention (retrieval, restudy, sleep, or wake), SIO = Sleep Inertia Offset / Retention Period, DR = Delayed Recognition.

The memory task was based on previous protocols which have employed retrieval training and sleep (Ferreira et al., 2019; Liu et al., 2021; Schreiner et al., 2018). Open Sesame v.3.3.10 software was used to create and run each phase of the memory task (Mathôt et al., 2012). The memory task began with a learning phase, then immediate recognition, one of four intervention conditions (retrieval, restudy, sleep, or wake), and a final delayed recognition. The intervention conditions across sessions were experienced in a pre-determined, counterbalanced order. This counterbalancing was random, except that the subjects always completed one wakeful condition before the sleep condition, to increase their familiarity with the environment and therefore increase their likelihood of napping. The paired associates used in the memory task consisted of randomly paired MST object pictures and auditory words. All subjects were presented with the same object-word pairs, in a randomised order for each condition by subject. A different set of object-word pairs were used in each of the 4 conditions. For the object in each object-word pair, there was an object in a different object-word pair that was its similar object from the MST. Objects and words in the same pair that were highly related to each other (e.g., dog-cat) were manually identified and reshuffled. A visualisation of the memory task can be found in Figure 2. The following sections detail the memory task protocol.

**Fig. 2.**
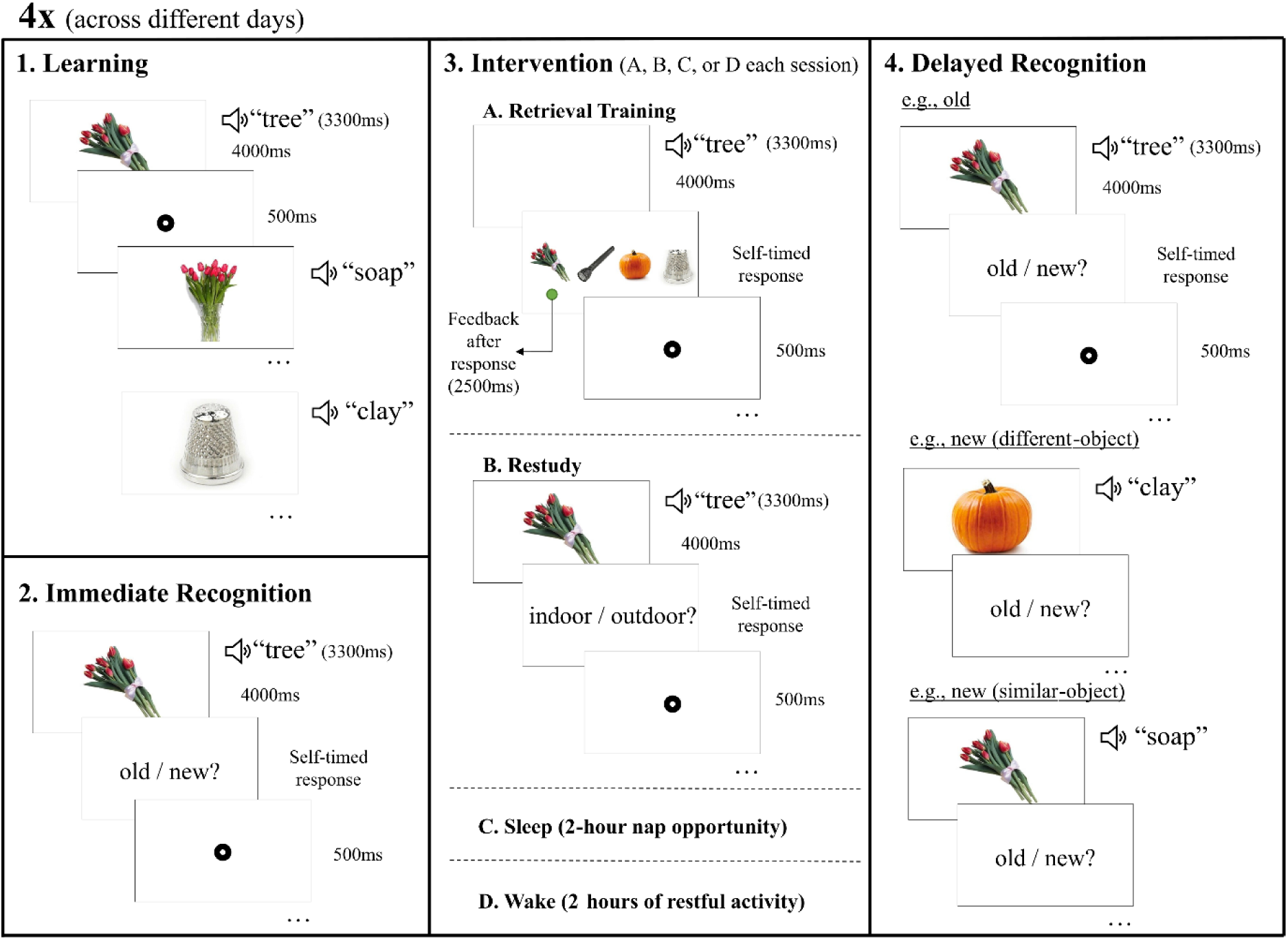
Memory task. 1. In the learning phase, subjects were exposed once to each of the object-word pairings. 2. Subjects then underwent immediate recognition testing, to distinguish between original (old) and rearranged (new) pairings, based on the pairings learnt during learning. An example of an old pairing is shown here. 3. Subjects underwent one of four intervention conditions. A. Involved retrieval training, where subjects heard each word, recalled the object, then picked it from distractors with feedback after. B. Subjects were exposed to all the pairs again and had to indicate if the object belonged indoors or outdoors. C. Subjects had a 2-hour nap opportunity. D. Subjects engaged in light activities. 4. Delayed recognition involved the same procedure as immediate recognition, where subjects distinguished between old and new pairings of the objects and words. This includes an example of an old pairing, new pairing with a different object to the object paired with the original word, and a new pairing with a similar object to the object paired with the original word.

#### 2.3.1. Learning

In each session, subjects learnt 104 object-word pairs. This involved viewing the image of an object on a white screen for 4000 ms and a sound of the paired auditory word was played 700 ms after the object onset. The images were displayed in colour, 400 pixels wide, in the centre of the screen. After each object was displayed, a fixation dot (8 pixels wide with a 2-pixel wide centre hole) was presented for 500 ms, before the next object was shown. Each pair was presented once.

#### 2.3.2. Immediate Recognition

Immediately following learning, recognition memory of the pairs was tested by exposing subjects to each object and word again in the same sequence as the learning phase. This was done using an old/new paradigm, where 72 object-word pairs were presented in their correct pairings (old pairs), and 32 in a rearranged pairing (new pairs). To create the rearranged pairings, pairs were shuffled by substituting the object in each pair with another object of a different pair (16 similar objects, 16 different objects). After each pair was presented, a new white screen displayed the prompt “old or new?” Subjects responded via keypress, based on whether they believed that the pairing was presented as it was originally learnt (old / “Q”), or if it was rearranged (new / “P”). Before starting the task, subjects were encouraged to respond quickly, but optimize accuracy over speed. After they responded to the prompt, a fixation dot appeared on the screen for 500 ms before the next object-word pair was shown. No feedback was given to subjects on recognition accuracy performance. To avoid introducing interference effects with the original encoding episodes, the 32 rearranged (new) pairs were not used in subsequent phases.

#### 2.3.3. Intervention

For each of their four visits, subjects completed one of the four interventions, depending on the condition order they were assigned. Each intervention period took 120 minutes and EEG was recorded, apart from the wake condition. After the intervention, all conditions included a 45-minute retention period. This also served as a sleep inertia offset period for the nap condition, to reduce the impact of post-sleep cognitive impairments on their memory performance (Hilditch & McHill, 2019). During this break, subjects remained in the laboratory and engaged in light activities (e.g., reading or scrolling on social media), simulating how they typically spend their free time. The KSS was completed at the end of the intervention. The four interventions are outlined below.

##### 2.3.3.1. Retrieval Training

Subjects completed six rounds of retrieval training practice with the initially learnt object-word pairs. The 72 object-word pairs were randomly divided into two equal length lists, with each list being presented three times. Within each round, the order of presentation for the object-word pairs was randomised per subject. Practice rounds took 10−12 minutes, with 8–10-minute breaks between so that the practicing was spaced out over the 120-minute period.

The retrieval training practice had subjects hear the word of each pair, recall the paired object from their memory, then pick the object that was originally paired with the word from distractors. Subjects were presented a blank white screen and heard the word of one of the object-word pairs that were previously learnt. Subjects were instructed to imagine the paired object on the screen, when they heard the word. After 4000 ms, four object images that were part of previously learnt pairs were displayed 120 pixels wide, equally spaced across the width of the screen. Of the four objects, one of them was the original object that the word was paired with, while the other three were objects from other pairs. The distractor options were pre-determined and consistent across all subjects, but they were different each of the three times that a word was practiced. For every single pair that was practiced, exactly one round included the original object’s similar MST object as one of the distractors. Subjects were instructed to choose the object that they believed they previously learnt as paired with the word they just heard. Each distractor object had a black letter displayed above it which indicated the corresponding response key (Q, E, I, or P) required to select the object below it. 500 ms after their response, subjects received feedback on their selection via a green dot (8 pixels wide with a 2-pixel wide centre hole) placed below the correct object for 2000 ms. Subjects were instructed to use this feedback to learn the pairings and improve their memory. After the feedback was presented, a fixation dot was shown in the middle of the screen for 500 ms, before the next word was played on a blank screen. For the breaks between each round, subjects engaged in light activities.

##### 2.3.3.2. Restudy

In the restudy condition, subjects followed the same six-round study-break structure as the retrieval training intervention, practicing 36 of the 72 pairs each round. Restudy practice involved being exposed to both constituents of each word pair again. For each object-word pair, subjects were presented the object and word exactly as was done during learning, in a subject-specific randomised order. An object image from each pair was presented in the centre of the screen for 4000 ms, followed by the originally paired word presented auditorily 700 ms later. To maintain subjects’ attention, after each pair was presented, a screen prompted subjects to decide if the object of that pair would be commonly found indoors or outdoors. This prompt was displayed in the centre of a white screen, with the words “Indoors / Outdoors?” with the letter “Q” and underneath the word “Indoors,” and the letter “P” underneath “Outdoors.” These letters underneath the question depicted the keypress required to select the corresponding option. Subjects were instructed to treat this interactivity task as secondary to their practicing of the pairs. After subjects responded, a fixation dot appeared for 500 ms before the next object-word pair was presented.

##### 2.3.3.3. Sleep

Subjects were given a 120-minute nap opportunity. Subjects slept in a quiet, darkened bedroom, and to promote sleep white noise was played at approximately 37db throughout the nap opportunity. The researcher remained in the adjacent room, monitoring the EEG signal for signs of sleep and any signal quality issues.

##### 2.3.3.4. Wake

In the wake condition, subjects were supervised while they engaged in light activities (e.g., reading a book or watching a movie) for 120 minutes, to replicate their typical use of free time. During this time, subjects’ EEG was not recorded, but they remained in the laboratory with the EEG cap on.

#### 2.3.4. Delayed Recognition

Similar to the immediate recognition phase, subjects were presented with the same old/new task, but with a new assignment of pairs as either old or new. Subjects were exposed to the 72 object-word pairs previously practiced, which were presented using the same sequencing as the immediate recognition phase. For half of the 72 pairs, the word of each pair was presented with its originally paired object from the learning phase. The other half (36) of the object-word pairs were novel (new) pairings. Half of the new pairings were similar-object lures, whereas the other half were different-object lures. Subjects were prompted, by the screen following the object, to indicate if the pairing was old or new, receiving no feedback on their accuracy. After they answered, a fixation dot appeared for 500 ms before the next pair was presented. Whilst the order of presentation of pairs was randomised for each subject, all shuffled pairings and assignments to old or new categories were pre-determined to remain consistent across all subjects. After the delayed recognition test, subjects had the EEG cap removed, and left the laboratory.

### 2.4. Data Analysis

#### 2.4.1. Behaviour Analyses

##### 2.4.1.1. Recognition Accuracy (percentage correct)

Recognition accuracy was calculated for each subject and condition during the old/new task at immediate and delayed recognition. During the old/new tasks, each word was presented with either its original paired-object (original), the MST similar-object to the original paired-object (similar), or a different object from another pair (different). To test recognition accuracy for each of these categories, scores were calculated per paired-object type used at delayed recognition. Accuracy was calculated by computing the percentage of correct response, i.e., selecting “old” when the pairing was presented as it was learnt originally (original objects), or selecting “new” when the object and word presented were from different pairs (similar and different objects). Therefore, higher recognition accuracy scores reflect greater ability to recognise the learnt associations between objects and words. Specifically for the similar lures, lower recognition accuracy reflects less veridical memory of the pairs. Values were determined to be outliers if they were more than 1.5 times the interquartile range above quartile 3, or below quartile 1. Outliers were removed for recognition accuracy scores, as well as every other continuous variable described hereafter.

##### 2.4.1.2. Behaviour Model

To test if retrieval training and sleep conditions lead to a greater propensity to endorse similar lures and enhanced accuracy for old pairs, we conducted a linear mixed effects model using R v4.3.0 (R Core Team, 2022) and the *lme4* package (Bates et al., 2015). The model aimed to predict delayed recognition accuracy scores, using intervention condition (i.e., retrieval, restudy, sleep, and wake) and paired-object type (i.e., original, similar, and different) as fixed effects. This model also controlled for immediate recognition accuracy and KSS scores. Subject ID was included as a random effect. Pre-stimulus activity and LPC amplitudes were scaled in this model, by subtracting the mean from all values. Treatment contrast coding was applied to determine significant differences between paired-object types within each condition.

#### 2.4.2. ERP Analyses

##### 2.4.2.1. EEG Preprocessing

The EEG signals were processed using MNE-Python v.1.7.0 (Gramfort et al., 2013). Activity from each intervention period was re-referenced to the mastoid electrodes M1 and M2 and down-sampled to 250 Hz. A notch filter (50 Hz) removed interference from the Australian mains power supply. Also, a zero-phase finite impulse response filter using a Hamming window was applied to remove slow signal drifts below 1 Hz and high frequency activity above 40 Hz. Then, an Independent Component Analysis was applied to the retrieval and restudy intervention’s data to correct electrooculographic (EOG) artifacts, excluding components which correlated the most strongly with EOG events (via the create_eog_epochs function in MNE-Python). EEG was segmented into epochs beginning –100 ms before each word onset, and ending 1000 ms after word onset. Lastly, the *AutoReject* package (Jas et al., 2017) was used to reject bad epochs and repair bad channels via topographic interpolation. Sleep scoring of EEG data during the sleep intervention was performed as per established procedures (Berry et al., 2017).

##### 2.4.2.2. Parietal Old/New Effect

A grand average was calculated across all subjects’ epochs, to visualise ERP differences between hit (correctly identified as “old”) and correct rejection (correctly identified as “new”) object-word pairings, averaged across left parietal electrodes (CP1, CP3, CP5, Pz, P1, P3, P5, P7, PO7). Another grand average was calculated across the same region to visualise false alarm (incorrectly endorsed as old) and correct rejection object-word pairings. This region of interest was informed by prior literature, which also informed the use of a 500–800 ms window post-word onset (Paller et al., 1995; Paller & Kutas, 1992; Rugg, 1995; Rugg & Curran, 2007; Smith, 1993).

We then aimed to determine if we had successfully detected the parietal old/new effect that the LPC amplitude was higher for hits than correct rejections, and if this was reduced for retrieval and sleep conditions. We averaged the LPC amplitude values across the 500–800 ms window and the left-parietal region of interest for hits (correctly identified as “old”) and correct rejections (correctly identified as “new”) pairs. We then conducted a linear mixed-effects model, predicting mean LPC amplitude values from the fixed effects of condition (i.e., retrieval, restudy, sleep, and wake) and score (hits and correct rejections). This model controlled for pre-stimulus activity (–100–0 ms), with subject ID as a random effect.

##### 2.4.2.3. False Alarm Modelling

We also aimed to capture episodic recollection to false alarms, using the parietal old/new effect on the LPC. Firstly, we aimed to see if we could detect a parietal old/new effect with false alarms (incorrectly identified as “old”) compared to correct rejections (correctly identified as “new”). We conducted a paired-samples *t*-test to determine if LPC amplitudes across the 500–800 ms window and the left-parietal region, significantly differed between false alarm and correct rejection responses. A baseline correction was applied by subtracting the pre-stimulus activity from the LPC amplitude. Cohen’s *d* was used to estimate effect size.

We then investigated LPC amplitudes for different types of false alarms. To isolate instances when subjects perceived that they remembered a pair, we only analysed the LPC windows belonging to three groups of paired-object-scores; hits to original objects (correctly identified as “old”), false alarms to similar objects, and false alarms to different objects (incorrectly identified as “old”). To ascertain the episodic recollection beyond what was experienced by the pairs correctly identified as “new” (i.e., the parietal old/new effect) we calculated LPC amplitude averages, per subject and paired-object-score, and subtracted each subject’s LPC amplitude mean for correctly identified new pairs. We then used this score in the modelling described below.

We conducted another linear mixed-effects model to determine if retrieval and sleep led to less episodic recollection for old endorsements of similar-object pairings. We used the mean LPC difference amplitudes as the outcomes for the model, predicted by condition (i.e., retrieval, restudy, sleep, and wake) and paired-object-score (i.e., original hits, similar false alarms, and different false alarms). This model controlled for pre-stimulus amplitude and used subject ID as a random effect. Pre-stimulus activity and LPC amplitudes were centred in this model, by subtracting the mean from all values. We then applied treatment contrast coding to determine if significant differences existed between every paired-object-score within each intervention condition.

## 3. Results

Table 1 displays subjects’ sleep characteristics during the sleep intervention. Every subject entered NREM sleep, four subjects experienced rapid eye movement (REM) sleep, and only five subjects did not enter SWS.

**Table 1.**
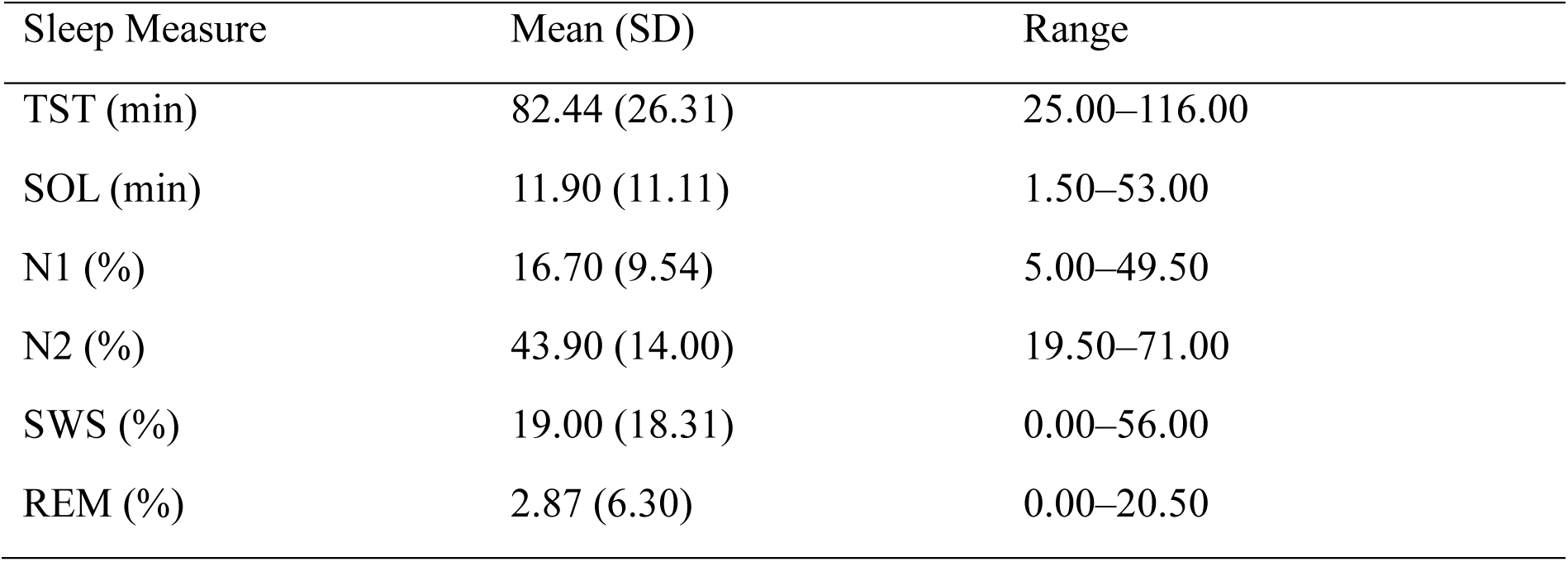
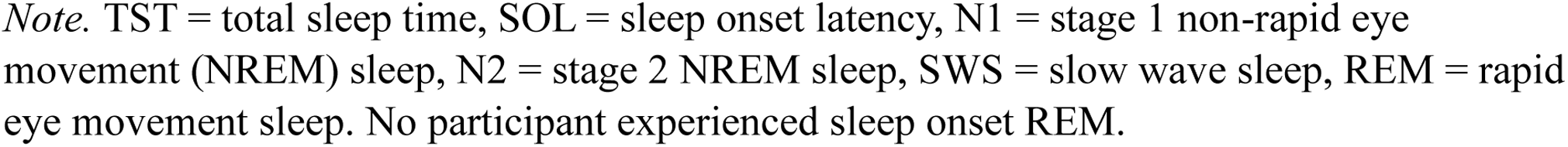
Sleep characteristics for the sleep intervention.

| Sleep Measure | Mean (SD) | Range |
| --- | --- | --- |
| TST (min) | 82.44 (26.31) | 25.00–116.00 |
| SOL (min) | 11.90 (11.11) | 1.50–53.00 |
| N1 (%) | 16.70 (9.54) | 5.00–49.50 |
| N2 (%) | 43.90 (14.00) | 19.50–71.00 |
| SWS (%) | 19.00 (18.31) | 0.00–56.00 |
| REM (%) | 2.87 (6.30) | 0.00–20.50 |

*Note.* TST = total sleep time, SOL = sleep onset latency, N1 = stage 1 non-rapid eye movement (NREM) sleep, N2 = stage 2 NREM sleep, SWS = slow wave sleep, REM = rapid eye movement sleep. No participant experienced sleep onset REM.

### 3.1. Retrieval training, but not sleep, simultaneously increased recognition accuracy and similar lure endorsement

The recognition accuracy scores for immediate and delayed recognition phases are displayed in Table 2, per condition and paired-object type.

**Table 2.** Recognition accuracy per paired-object type in the old/new task.

| Paired-Object | Condition | Immediate Recognition<br>Accuracy (%) | Delayed Recognition<br>Accuracy (%) |
| --- | --- | --- | --- |
| Original | Retrieval | 66.30 (16.85) | 92.62 (8.23) |
|  | Restudy | 67.24 (16.86) | 86.89 (14.91) |
|  | Sleep | 67.01 (14.02) | 70.25 (16.90) |
|  | Wake | 63.59 (13.52) | 66.67 (14.80) |
| Similar | Retrieval | 60.26 (21.16) | 71.55 (16.43) |
|  | Restudy | 59.13 (19.53) | 76.38 (15.42) |
|  | Sleep | 59.10 (19.01) | 69.19 (18.15) |
|  | Wake | 59.72 (16.75) | 55.76 (18.00) |
| Different | Retrieval | 77.06 (18.53) | 92.83 (13.47) |
|  | Restudy | 78.00 (19.94) | 88.22 (20.12) |
|  | Sleep | 74.34 (19.30) | 80.28 (18.29) |
|  | Wake | 68.06 (18.13) | 79.42 (17.89) |

*Note.* Means are reported, with standard deviations in brackets. Recognition accuracy refers to subjects' percentage of correct answers in an old/new task.

Our first linear mixed-effects model tested the hypothesis that retrieval and sleep would lead to both a greater accuracy in identifying original objects and greater endorsement of similar paired-object lures. This model revealed that delayed recognition accuracy was significantly predicted by an interaction between intervention condition (i.e., retrieval, restudy, sleep, and wake) and paired-object type (i.e., original, similar, and different). Table 3 depicts the results for each main and interaction effect in the model. Figure 3 visualises the interaction between condition and paired-object type to predict delayed recognition accuracy, showing that retrieval led to a greater accuracy for identifying original pairs, but a far lower accuracy for identifying similar object lures.

**Table 3.** Main and interaction effects for the behaviour model.

| Effect | *df* | $\chi^2$ | *p* |
| --- | --- | --- | --- |
| Condition | 3 | 92.90 | <.001 |
| Paired-object | 2 | 35.73 | <.001 |
| KSS | 1 | 0.04 | 0.845 |
| Immediate recognition accuracy | 1 | 148.60 | <.001 |
| Condition * paired-object | 6 | 39.14 | <.001 |

Note. *df* = degrees of freedom.

**Fig. 3.**
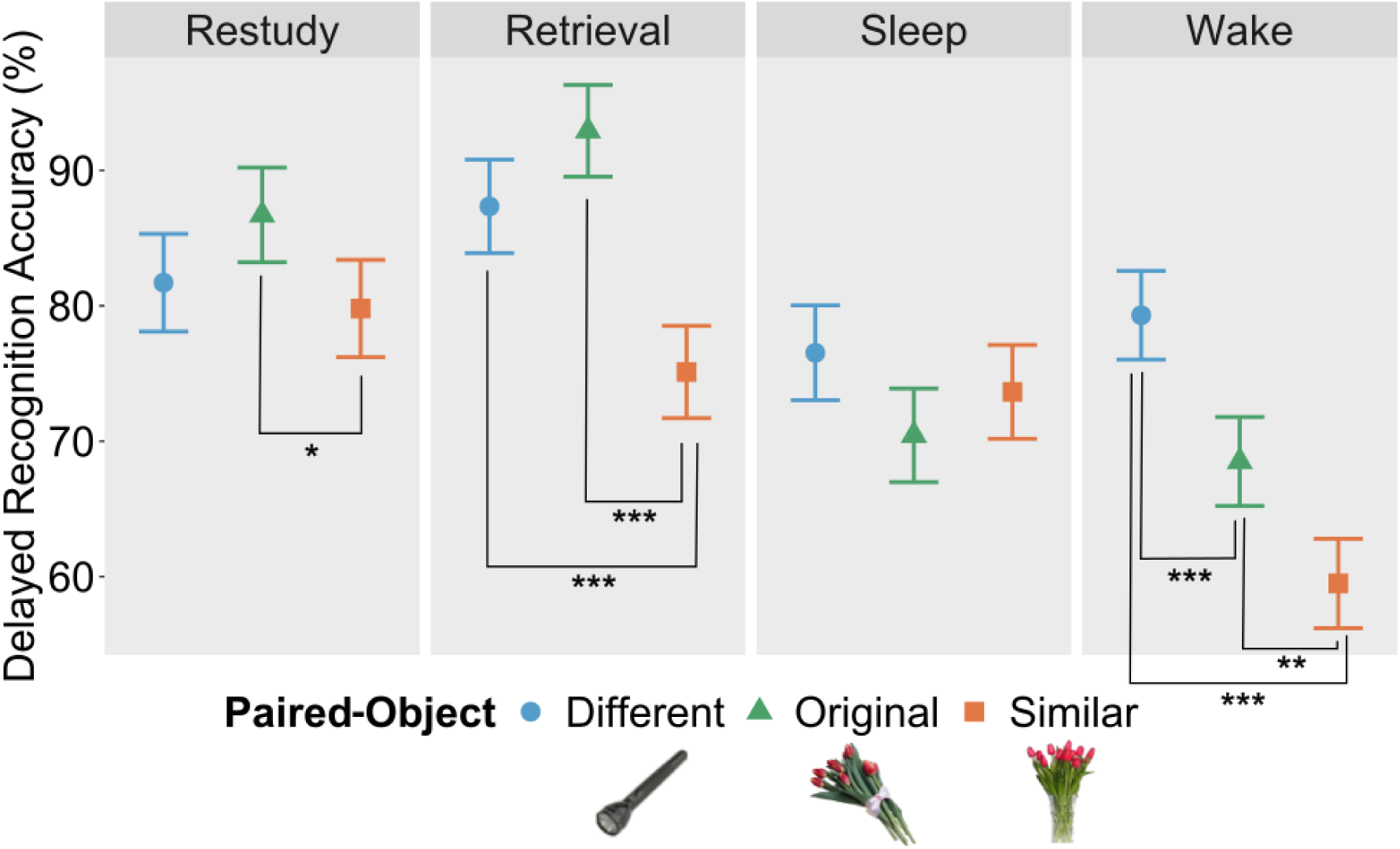
Modelled delayed recognition accuracy per intervention condition and paired-object type. The x-axis represents the type of paired-object to the word, the pairing of which was evaluated by subjects as old or new. The y-axis represents subjects’ recognition accuracy for each paired-object type. The facets depict the results of each intervention condition. The error bars represent the 83% confidence intervals. Significant treatment contrasts are marked as: \**p*<.05, \*\**p*<.01, and \*\*\**p*<.001.

### 3.2. LPC amplitudes for false endorsements of similar-object lures were lower than that of different-object lures, only after retrieval training

To determine if the parietal old/new effect was reduced for the sleep and retrieval conditions, we performed a linear-mixed effects model predicting LPC window mean amplitudes from intervention condition (retrieval, restudy, sleep, and wake) and score (hits and correct rejections). The test revealed a significantly greater LPC amplitude for old pairs compared to new pairs, demonstrated the parietal old/new effect. However, the model did not detect a difference in the parietal old/new effect between conditions. The model results are reported in Table 4. The grand average plot (Figure 4) depicts the larger amplitude for hits over correct rejections, approximately 500–800 ms post-word onset.

**Table 4.** Main and interaction effects for the parietal old/new model.

| Effect | *df* | $\chi^2$ | *p* |
| --- | --- | --- | --- |
| Condition | 3 | 2.87 | .412 |
| Score | 1 | 27.65 | <.001 |
| Pre-stimulus amplitude | 1 | 23.36 | <.001 |
| Condition * score | 3 | 2.10 | .552 |

Note. *df* = degrees of freedom.

**Fig. 4.**
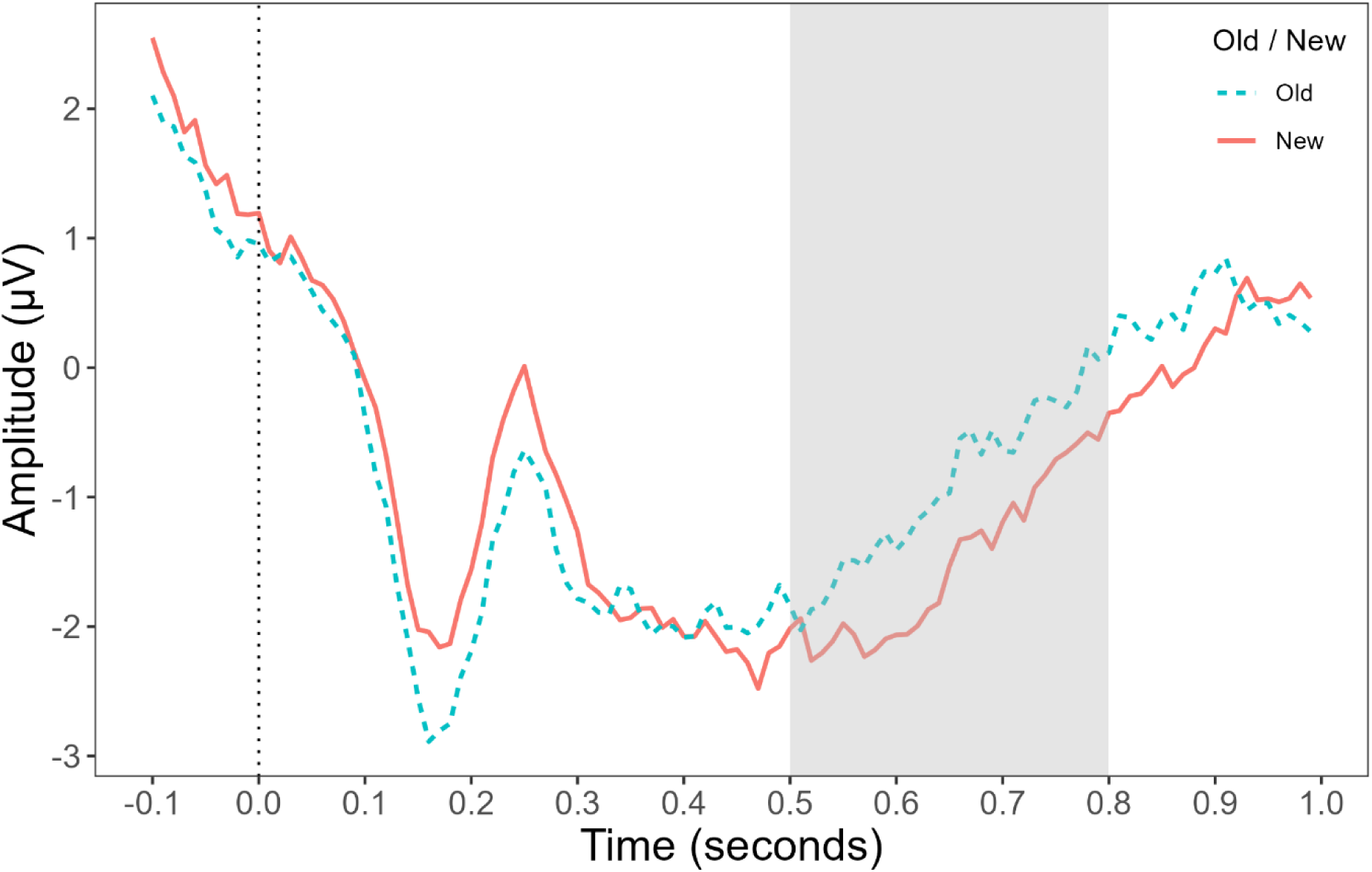
Grand average plot for hits and correct rejections post-word onset. The plot depicts the grand average for correct responses across left parietal electrodes. The x-axis is the time in seconds, relative to the word onset. A vertical dotted line marks the word onset time. The grey column highlights the late positive component window from 500–800 ms post-word onset. The y-axis is the amplitude of the epochs in microvolts. The amplitude for old pairs (hits) are depicted in blue with a dashed line, new pairs (correct rejections) are depicted in red with a solid line.

To investigate the LPC amplitude for false alarms, we first plotted the grand average ERP with false alarms and correct rejections, in Figure 5. A paired-samples *t*-test revealed a significantly higher amplitude for false alarms than correct rejections with a small effect size, *t*(102)=-2.46, *p=*.016, *d*=-0.24.

**Fig. 5.**
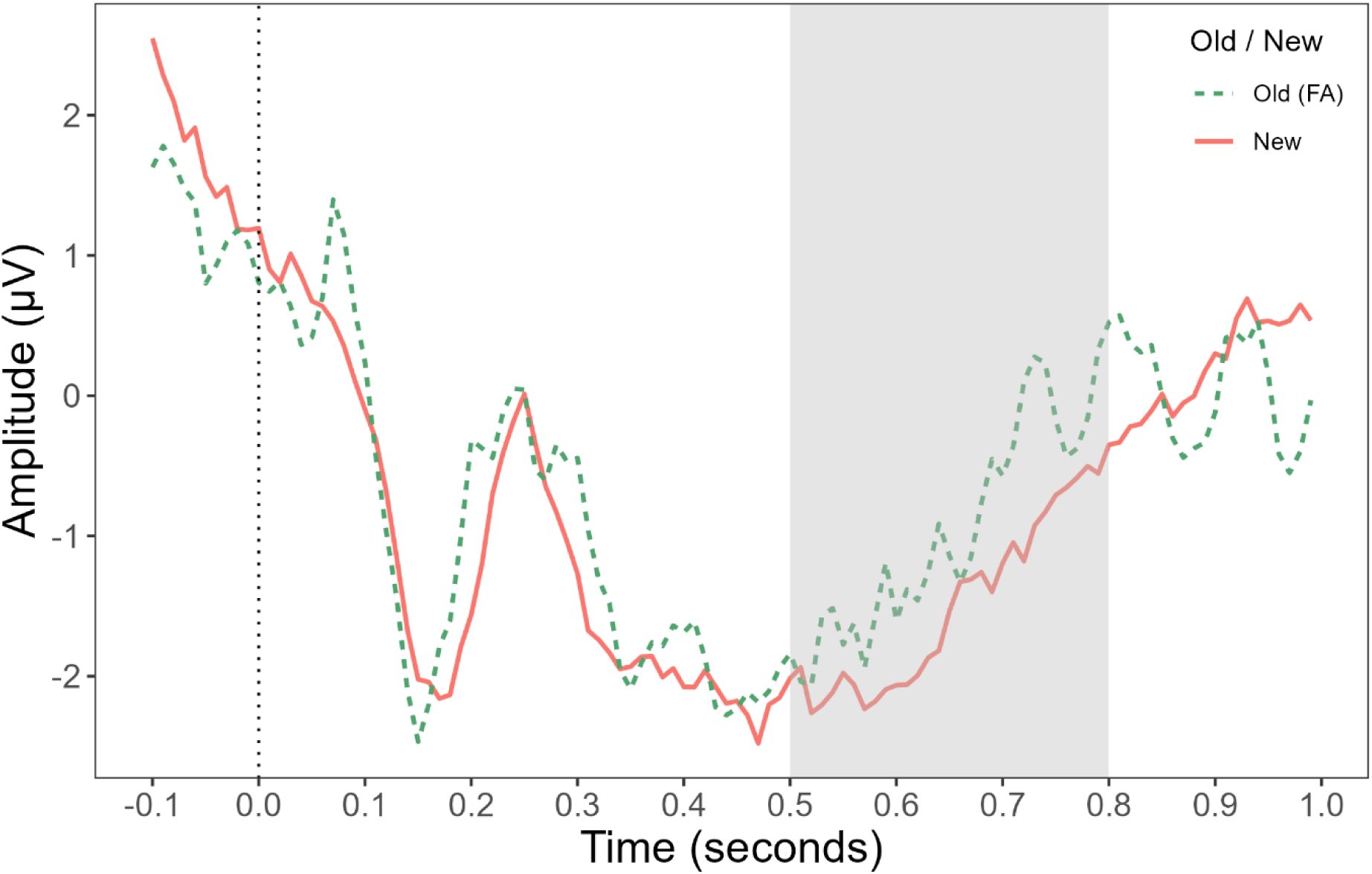
Grand average plot for false alarms and correct rejections post-word onset. The plot depicts the grand average for false alarms across left parietal electrodes. The x-axis is the time in seconds, relative to the word onset. A vertical dotted line marks the word onset time. The grey column highlights the late positive component window from 500–800 ms post-word onset. The y-axis is the amplitude of the epochs in microvolts. The amplitude for old pairs (false alarms) are depicted in green with a dashed line, new pairs (correct rejections) are depicted in red with a solid line. FA = false alarms.

Following this grand average, we calculated the difference between LPC amplitudes for each paired-object-score (i.e., original object hits, similar object false alarms, and different object false alarms) against LPC amplitudes for correct rejections. A linear mixed-effects model revealed that these LPC difference amplitudes were significantly predicted by an interaction between intervention condition (i.e., retrieval, restudy, sleep, and wake) and paired-object-score (i.e., original hit, similar false alarm, and different false alarm). All interaction and main effects in the model are depicted in Table 5. Figure 6 depicts that sleep and wake conditions had indistinguishable LPC amplitudes for each paired-object types. The retrieval training condition had lower LPC amplitudes for similar compared to different paired-object types, but restudy showed the opposite pattern.

**Table 5.**
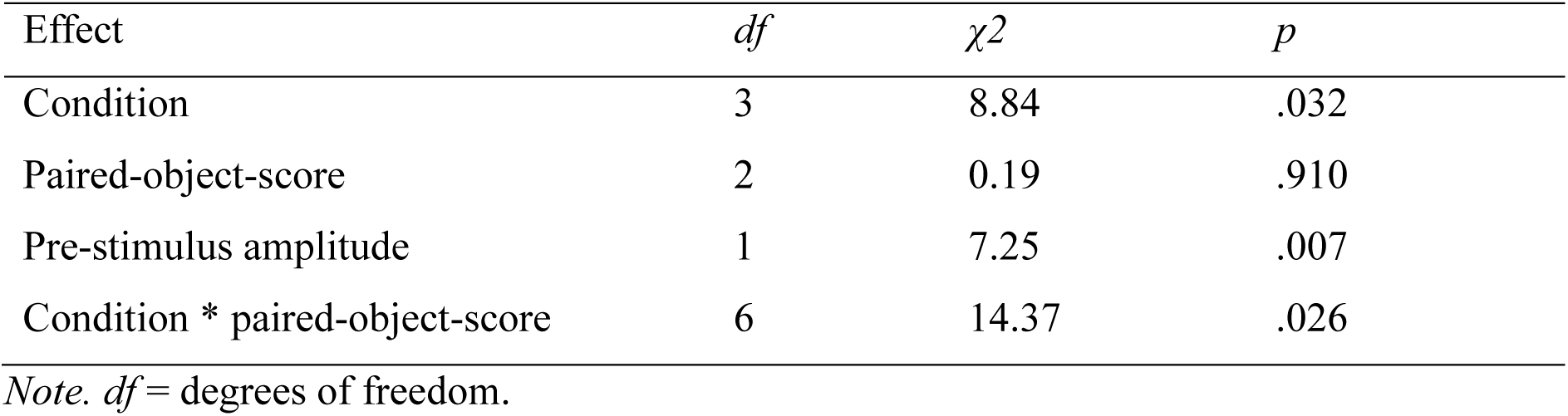
Main and interaction effects for the paired-object-score model.

**Fig. 6.**
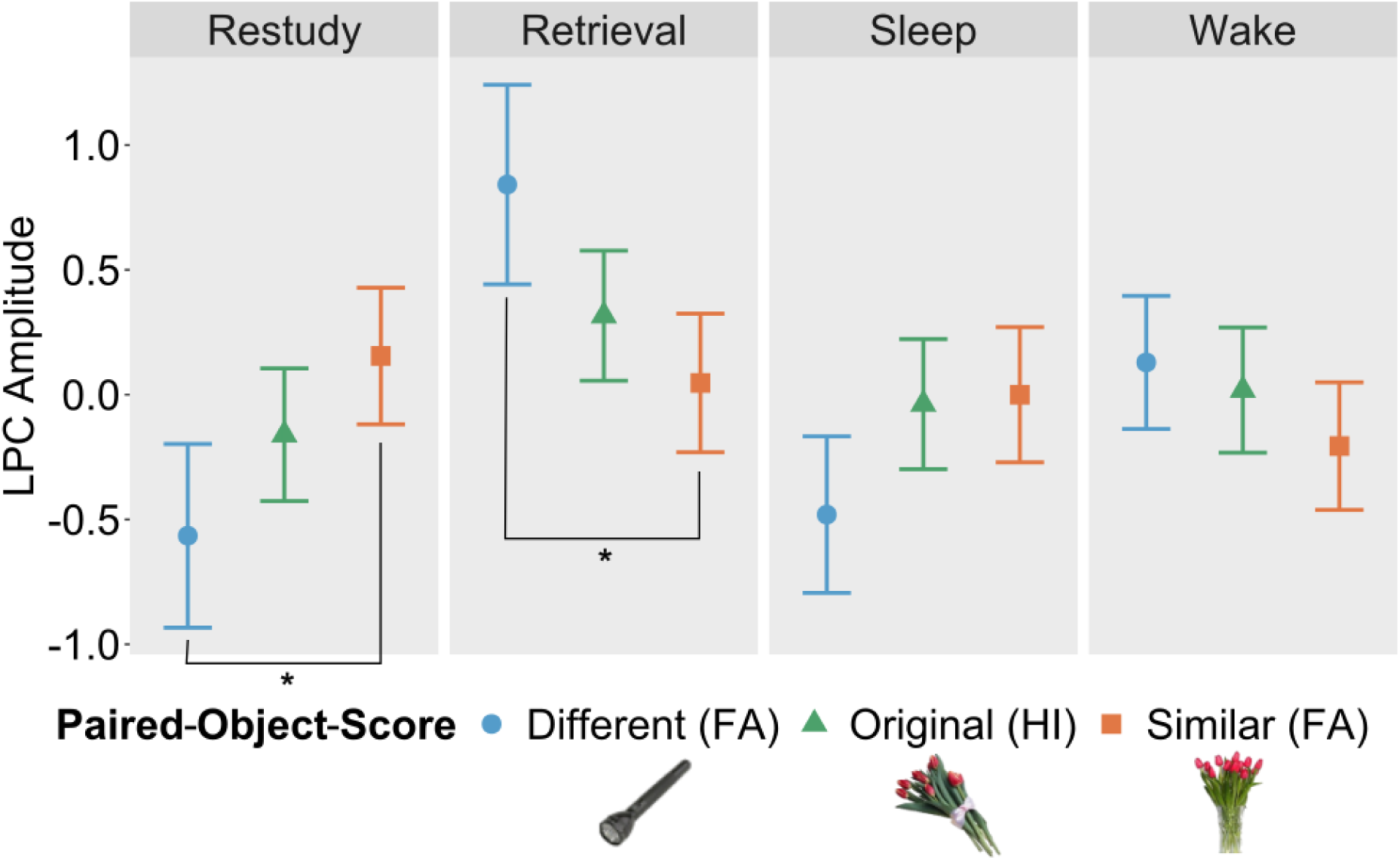
Modelled LPC amplitude per intervention condition and paired-object-score. The x-axis represents the type of paired-object-score, which is a combination of the paired-object to the words, and the accuracy of evaluating the pair as old. FA= false alarm (i.e., subject answered “old” to a “new” pairing), HI=hit (i.e., subject answered “old” to an “old” pairing). The y-axis represents the modelled and centred late positive component (LPC) amplitude for each paired-object-score, minus the amplitude for correct rejections. The facets depict the results of each intervention condition. The error bars represent the 83% confidence intervals. Significant treatment contrasts are marked as: \**p*<.05, \*\**p*<.01, and \*\*\**p*<.001.

### 4. Discussion

This study aimed to address if sleep-based and retrieval-mediated memory consolidation lead to comparable endorsements and neural correlates of different stimuli pairings. We found that retrieval training led to higher overall recognition accuracy, but a greater endorsement of similar-object lures. Conversely, recognition accuracy after sleep was relatively equivalent across different object lures and hit accuracy. Although we detected a parietal old/new effect on the LPC, indexing recollective processes, the magnitude of this effect was not modulated by intervention condition. Instead, there were differences between the intervention conditions regarding the LPC amplitudes for false alarms to similar- and different-object lures. Sleep and wake demonstrated no differences between any paired-object-score type, whereas retrieval resulted in higher recollection for different-object lures than similar-object lures, and restudy demonstrated the opposite. In conjunction, the behavioural and neural findings can speak to the different effects of the interventions on the recollection of memories.

Every condition, except sleep, led to poorer discrimination for similar lures than for old pairs and different lures. This difference was the most pronounced for the retrieval training condition, which demonstrated the best overall accuracy, but a comparatively much lower rate of similar-object lure discrimination. This may result from retrieval training promoting the fast semanticization of memory traces, which would have enhanced memory accuracy for old pairs and different-lures through those pairs being incorporated into sparse neocortical stores (Antony et al., 2017). This may have been what led to more false alarms when faced with semantically similar lures, as their discrimination hinged primarily on the contextual features that are less prioritised during neocortical integration (McClelland et al., 1995; Winocur & Moscovitch, 2011). Contrastingly, similar-lure discrimination was highest in the restudy condition, supporting the predictions that restudy reinstates strong episodic traces of memories (Antony et al., 2017). Contrary to our predictions, sleep did not lead to worse similar-lure discrimination, but instead a comparable ability to distinguish similar-lures, different-lures, and recognise old stimuli. Instead, every other condition that involved some wakeful interference was worse at similar-lure discrimination than other measures of memory accuracy. This could speak to sleep maintaining both the semantic and episodic traces of memories, and may further reflect a task-relevant selective maintenance of episodic features that were required for lure discrimination. This is in line with the roles of sleep in saving memories from short-term decay (Denis et al., 2023; Dumay, 2016; Rasch & Born, 2013), enhancing memory accuracy overall, and demonstrating task dependencies (Cordi & Rasch, 2021; Diekelmann et al., 2009). Further insights into what underscores these behavioural accuracy differences can be explored via the LPC and episodic recollection experienced.

Our results demonstrated a clear parietal old/new effect on the LPC, whereby amplitudes were higher for hits versus correct rejections. This is in line with studies that have detected this effect across numerous paradigms (Duzel et al., 1997; Paller et al., 1995; Paller & Kutas, 1992; Rugg, 1995; Rugg & Curran, 2007; Vilberg et al., 2006; Wilding, 2000; Xie et al., 2024; Yick & Wilding, 2008). Despite our predictions, we did not detect any condition difference in the magnitude of the parietal old/new effect. This is surprising, as the effect reliably indexes episodic recollection, which sleep and retrieval have been shown to influence in previous studies (Gao et al., 2016; Mograss et al., 2008; Palmer et al., 2013; Rosburg et al., 2014; Sweegers et al., 2015; Zeng et al., 2021). One key difference in our design that may explain this is that our pair constituents (objects and words) in the old/new tasks were technically all “old” as they were all learnt in pairs before, but it was the ways that their pairings were intact or rearranged that were “old” or “new,” respectively. Nie and Wu (2023) found that whilst the LPC amplitude for rearranged pairs were lower than intact pairs, it was still higher than that of new objects, as there was still recollection involved for their rearranged components. In our study, measuring this reduced difference between rearranged and intact pairs could have reduced our parietal old/new effect to a degree where condition differences were not statistically detectable.

We were able to detect differences in the parietal old/new effect between retrieval and restudy conditions, when accounting for false alarms to similar- and different-object lures. After retrieval training, the LPC amplitude was higher for different-object false alarms than similar-object false alarms, whereas restudy had a higher amplitude for similar- than different-object false alarms. The lower episodic recollection for similar-object false alarms aligns with predictions that retrieval training induces semanticization and greater dependence on semantic neocortical stores for memory recall (Antony et al., 2017; Brodt et al., 2016; Lifanov et al., 2021; Marin-Garcia et al., 2021; Ritvo et al., 2019). This idea also aligns with the greater behavioural generalisation we detected in the retrieval training condition. Alternatively, the higher recollection for different-object false alarms could have been brought about by the failure of retrieval training to create a gist-representation to base an accurate judgement on, with subjects then resorting to a less effective greater recollective effort to previous study episodes. This is supported by Trace Transformation Theory, which suggests that the semantic components of memories are preferably used for recollection after consolidation, and the hippocampus only engages when its episodic traces are required to support the process (Winocur & Moscovitch, 2011). For restudy, the lower recollection for different-object false alarms could have been due to failures to encode episodic details, leading to a false judgement. Moreover, the higher episodic recollection for similar-object false alarms could have occurred as this discrimination was still based on episodic memory components, but they did not have enough available for accurate discrimination. This is supported by the amplitude for similar-object false alarms being similar after restudy and retrieval training. Together, this is in line with restudy not inducing consolidation, and pre-consolidated memories being hippocampally dependent for memory recall (Antony et al., 2017; McClelland, 2013; McClelland et al., 1995).

For both the sleep and wake groups, there were no differences between the LPC amplitudes, for original objects and either lure type. Therefore, not engaging in practice did not impact the episodic recollection involved in hits versus false alarms of similar- and different-object lures. This is not in line with Jano et al.’s (2021) finding that sleep led to lower LPC amplitudes for false memories for semantically similar lures. Instead, our combination of no evidence for lower episodic recollection after sleep compared to wake, no differences in the paired-object-score types, and no behavioural indication of generalisation across sleep, indicates that we did not find evidence for gist-extraction in the sleep group.

This is surprising, given that research has found evidence for gist-extraction across sleep due to repeated reactivations in NREM (Born & Wilhelm, 2012; Cordi & Rasch, 2021; Gibson et al., 2022). Some research has shown that REM sleep may also be important for gist-extraction (Liu et al., 2024; Tononi & Cirelli, 2006); however we were not able to investigate this in our study as only a few participants experienced a small amount of REM. Further research should compare memory across retrieval training to sleep across the early (more NREM) and late (more REM) conditions of a split-night study, to see which stages create similar memory outcomes to retrieval training. Additionally, Jano et al. (2021) used the Deese-Roediger-McDermott paradigm, which, unlike our study’s task, is specifically designed for associative generalisation. Therefore, our difference in results could further support that sleep-based consolidation demonstrates task dependencies (Cordi & Rasch, 2021; Diekelmann et al., 2009), and may not extract gist unless more specifically probed to by the paradigm’s design.

In our study, the LPC amplitudes for similar-object lures were statistically indistinguishable between all conditions. However, in the context of the rest of the paired-object-score amplitudes within a condition, these amplitudes support evidence for how each condition shapes memories. It is also difficult for to disentangle whether a reduced parietal old/new effect means that a memory was not encoded well, or whether it is evidence for gist-extraction. This is why our interpretations of the parietal old/new effect were always made in conjunction with the relevant behavioural scores, to provide a comprehensive view of how the memories were experienced. Also, keeping comparisons between paired-object-score types mostly within conditions, meant that our interpretations were always about the proportional values experienced, rather than comparing amplitude values across different contexts.

Overall, these comparisons between the sleep and wake condition, and the restudy and retrieval condition, highlight key differences between consolidation across retrieval training and sleep. In this paradigm, sleep-based consolidation seemed to equalize episodic recollection for hits and different types of false alarms, as well as the accuracy for all different types of paired-object discrimination. This could speak to a role of sleep in selectively maintaining and enhancing task-relevant aspects of memories where similar-lure discrimination is a task-relevant goal. However, retrieval demonstrated evidence for generalisation across memories and gist-extraction, regardless of the task-relevant goal to maintain episodic details for similar-lure discrimination. Whilst gist-extraction across retrieval training improved overall memory in most cases, in the few cases it did not, recollection had to fall on the less-maintained episodic traces. This difference between sleep and retrieval training conditions could be due to the role of a nap compared to longer sleep periods; however, naps have been shown to demonstrate comparable NREM-specific memory effects to longer sleep periods (Chen et al., 2024; Kumral et al., 2023). Therefore, sleep may be able to act on memories differently based on goals, potentially through prior tagging of relevant memories (Stickgold, 2013). Instead, retrieval training may mimic some sleep-based consolidation findings through quick and consistent semanticization (Ritvo et al., 2019), that may not be as paradigm-specific as sleep (Cordi & Rasch, 2021; Diekelmann et al., 2009). These results speak directly to the need to compare sleep and retrieval training in the same paradigm to draw conclusions about their similarities.

Our research has uncovered various questions that future research should address to continue building a comprehensive view of memory consolidation across retrieval training and sleep. Our research isolated episodic recollection after the word presentation, as this was when all the information had been provided to subjects to make their old/new distinction. However, subjects were not explicitly instructed on what information to recollect during this distinction (i.e., the object, the word, the entire pair, or the association), so that their recollection could be better influenced by the intervention. Therefore, we cannot be certain what aspects of the memory were reflected in the episodic recollection we measured, and if the content differed across intervention conditions. This could be addressed by pattern recognition algorithms that can identify the reactivation patterns of memories that re-emerge within these windows, which would further reveal what aspects of these memories that each intervention prioritises for subsequent recollections. Additionally, theories of representational change across memory consolidation typically mention the changes within memories, as well between memories (Antony et al., 2017; McClelland et al., 1995; Ritvo et al., 2019; Winocur & Moscovitch, 2011). Our study addressed the within-memory changes from episodic to semantic dependence, so future research should investigate if between-item changes across sleep and retrieval training also demonstrate evidence for memory generalisation through inter-item representational similarities (e.g., between similar and different items). Lastly, our study demonstrated how false alarms to similar-object (semantic) lures differed from false alarms to different-object lures. To further demonstrate how episodic versus semantic details are maintained and depended on across retrieval training and sleep, future studies should compare the recollection involved in false alarms to both semantic and episodic lures.

In summary, this research found that sleep-based and retrieval-mediated memory consolidation created differences in behavioural and neural markers for generalisation, across equivalent consolidation opportunities. Whilst retrieval training induced a fast route to behavioural and neural semanticization, sleep demonstrated a task-relevant selective maintenance of episodic discrimination and recollection. These findings challenge existing theories that memory consolidation acts similarly across both interventions, and may imply that memory transformations during sleep are supplemented by processes that retrieval training cannot reliably replicate across wake.

## Acknowledgements

We thank Annaliese Anesbury and Maya Al Safadi for their assistance during data collection. We also thank the subjects for their time.

## CRediT authorship contribution statement

Hayley B. Caldwell: Conceptualization, Data curation, Formal analysis, Investigation, Methodology, Project administration, Validation, Visualization, Writing – original draft, Writing – review and editing.

Kurt Lushington: Conceptualization, Methodology, Project administration, Resources, Supervision, Writing – review and editing.

Alex Charburn: Conceptualization, Methodology, Project administration, Resources, Supervision, Writing – review and editing.

